# Lymphatic Thrombosis and Impaired Lymph Drainage in Cigarette Smoke-Associated Emphysema

**DOI:** 10.1101/2021.11.23.469393

**Authors:** Barbara D. Summers, Kihwan Kim, Zohaib Khan, Sangeetha Thangaswamy, Cristina C. Clement, Jacob McCright, Katharina Maisel, Sofia Zamora, Stephanie Quintero, Alexandra C. Racanelli, Jisheng Yang, Jeanine D’Armiento, Laurel Monticelli, Mark L. Kahn, Augustine M. K. Choi, Laura Santambrogio, Hasina Outtz Reed

## Abstract

The lymphatic vasculature is critical for lung function, but defects in lymphatic function in the pathogenesis of lung disease is understudied. In mice, lymphatic dysfunction alone is sufficient to cause lung injury that resembles human emphysema. Whether lymphatic function is disrupted in cigarette smoke (CS)-induced emphysema is unknown. In this study, we investigated lung lymphatic function in the pathogenesis of CS-induced emphysema. Analysis of human lung tissue revealed significant lung lymphatic thrombosis in patients with emphysema compared to control smokers that increased with disease severity. *In vitro* assays demonstrated a direct effect of CS on lymphatic endothelial cell integrity. In a mouse model, CS exposure led to lung lymphatic thrombosis, decreased lymphatic drainage, and impaired leukocyte trafficking that preceded emphysema. Proteomic analysis of lymph confirmed upregulation of coagulation and inflammatory pathways in the lymphatics of CS-exposed mice compared to control mice. These data suggest that CS exposure results in lung lymphatic dysfunction with thrombosis, impaired leukocyte trafficking, and changes in the composition of lymph. In patients with emphysema, lung lymphatic thrombosis is seen with increasing disease severity. These studies for the first time demonstrate lung lymphatic dysfunction after cigarette smoke exposure and suggest a novel component in the pathogenesis of emphysema.

## Introduction

Chronic Obstructive Pulmonary Disease (COPD) includes emphysema and chronic bronchitis and is commonly caused by cigarette smoke (CS). Despite extensive knowledge about the pathologic changes in the lung epithelium, blood endothelium, and the cellular mechanisms for lung injury in the pathogenesis of COPD, the lung lymphatic vasculature has not been well evaluated. The lung lymphatics drain fluid and traffic immune cells in the form of lymph from the lung parenchyma to the draining lymph nodes^1,2^. Though previously thought to be a passive conduit for lymph, an active role for the lymphatics in the inflammatory response has been increasingly the subject of investigation. Accordingly, defects in lymphatic function may play a role in the pathogenesis of disease, especially in the lungs, which are particularly dependent on lymphatic function due to the vulnerability of this organ to edema and its constant exposure to pathogens^3^.

We have previously shown that mice with lymphatic dysfunction develop lung tertiary lymphoid organs (TLOs) and lung injury with many features of human emphysema including hypoxia, breakdown of elastin, and increased MMP-12 expression^4^. TLOs are intricately associated with lymphatic vessels and resemble lymph nodes in their cellular organization and structure^5^. They are a common occurrence in lung injury and inflammation, including in COPD, where they may also be associated with disease severity^5-11^. Though lymphatic dysfunction is sufficient to cause TLO formation and lung injury that resembles emphysema in mice, it is not yet clear whether the TLOs that are seen in emphysema are associated with lymphatic dysfunction. In this study, we sought to uncover whether and to what extent lung lymphatic function is altered in the pathogenesis of emphysema and the mechanism by which this occurs.

Here, we report that lung lymphatic vessel thrombosis is increased in emphysema compared to control smokers and is associated with severe disease. Using a mouse model, we found that tissue destruction and emphysema alone was not sufficient to cause lung lymphatic thrombosis. However, lung lymphatic thrombosis and dysfunction was associated with injury to the lymphatic endothelium and changes in the composition of lymph after CS exposure which occur before the development of emphysema in mice. These data are the first to show a direct effect of CS on lung lymphatic function and to demonstrate lymphatic thrombosis in patients with emphysema.

## Results

### Lung Lymphatic Thrombosis in Severe Emphysema

To explore changes in lung lymphatics in the setting of CS-induced emphysema, we analyzed the lymphatics in lung tissue from patients with a history of cigarette smoking that have been and clinically and radiographically identified as having emphysema. Using immunohistochemical analysis for the lymphatic marker Podoplanin, we found no change in the density of lung lymphatics in patients with emphysema compared to control smokers (Figure 1A,B,I). This contrasts with previous studies that reported increased lung lymphatic density in human COPD^12,13^, though this may be due to differences in the lymphatic markers used and the area of the lung that was assessed. Close analysis revealed fibrin-rich thrombi in the lymphatics of patients with emphysema, in some cases obstructing the entire lumen of the vessel (Figure 1C-F). Lymphatic thrombosis was significantly increased in patients with very severe emphysema compared to moderate emphysema or control smokers (Figure 1J). TLOs were similarly increased in severe emphysema (Figure 1G,H,O), as seen in previous studies^6^. Interestingly, we identified thrombosed lymphatic vessels that were spatially associated with TLOs in these samples (Figure 1K-N).

**Figure 1:**
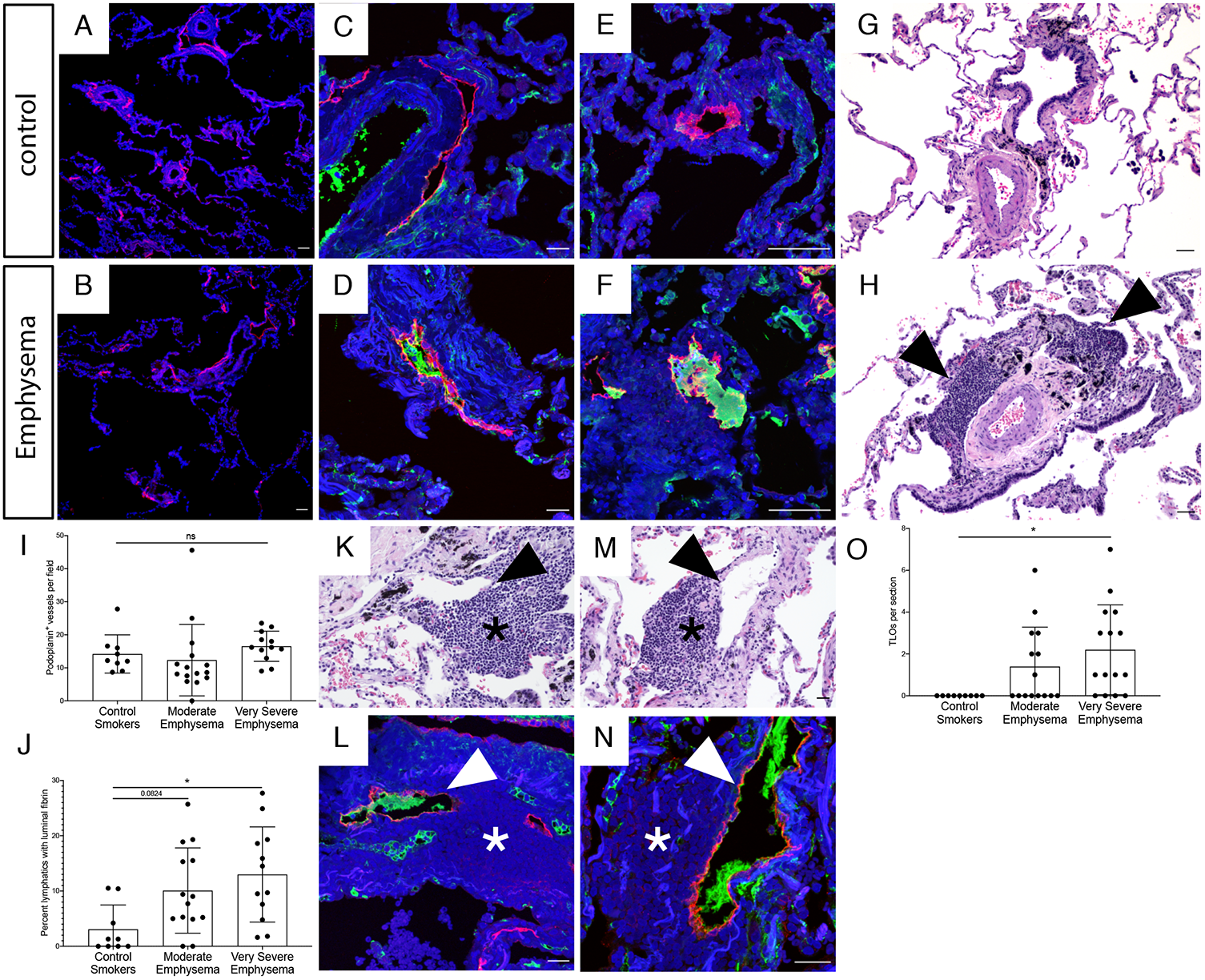
Lymphatic thrombosis in human emphysema. (A,B) Representative fluorescent immunohistochemical analysis of human lung tissue from patients with emphysema or control smokers for the lymphatic marker podoplanin (red). (C-F) Immunohistochemical staining for lymphatic thrombosis using podoplanin (red) and fibrinogen (green) in lung tissue from patients with emphysema (D,F) and control smokers (C,E). (G,H) H&E staining of lung tissue revealed TLOs in emphysema (H, arrowheads) but not control lung tissue (G). (I) Quantification of podoplanin^+^ lung lymphatics in tissue from control smokers and patients with moderate and very severe emphysema. (J) Quantification of podoplanin^+^ lymphatic vessels with luminal fibrin. (K-N) Serial H&E and immunohistochemistry serial sections (K,L and M,N) of lung tissue from patients with very severe emphysema demonstrating lymphatic thrombosis (arrowheads) spatially associated with TLOs (asterisks). (O) Quantification of TLOs in human lung tissue. Quantification lung lymphatic vessels was performed by counting podoplanin^+^ lymphatics in 10x images of lung tissue sections. At least 3 10x images were used for each patient tissue sample and the average lymphatic number was determined. Lymphatic thrombosis was quantified as the percentage of podoplanin^+^ lung lymphatics with luminal fibrin in each 10x image of the tissue section and was averaged from at least 3 10x images. TLOs were quantified as the total number of TLOs visualized in each lung tissue sample using H&E staining. All values are means ± SEM. *P* value calculated by ANOVA. \**P* <0.05. ns = not significant. Scale bars = 25µm.

### Lung Lymphatic Thrombosis Does Not Occur after Elastase-induced Emphysema in Mice

To expand on the finding of lung lymphatic thrombosis in human emphysema, we used a mouse model to ask whether alveolar destruction in emphysema itself results in lymphatic thrombosis. To do this, we used the elastase model, in which tracheal instillation of porcine pancreatic elastase results in severe lung injury and progressive alveolar breakdown resulting in emphysema within 21 days^14^. Elastase treatment did not affect lung lymphatic density (Figure 2A). In addition, we did not observe any significant lymphatic thrombosis in this model, despite severe emphysema and alveolar damage (Figure 2B-F). Interestingly, elastase treatment also did not result in TLO formation in association with the development of emphysema, even in areas with most significant alveolar enlargement and tissue destruction (Figure 2G-H).

**Figure 2:**
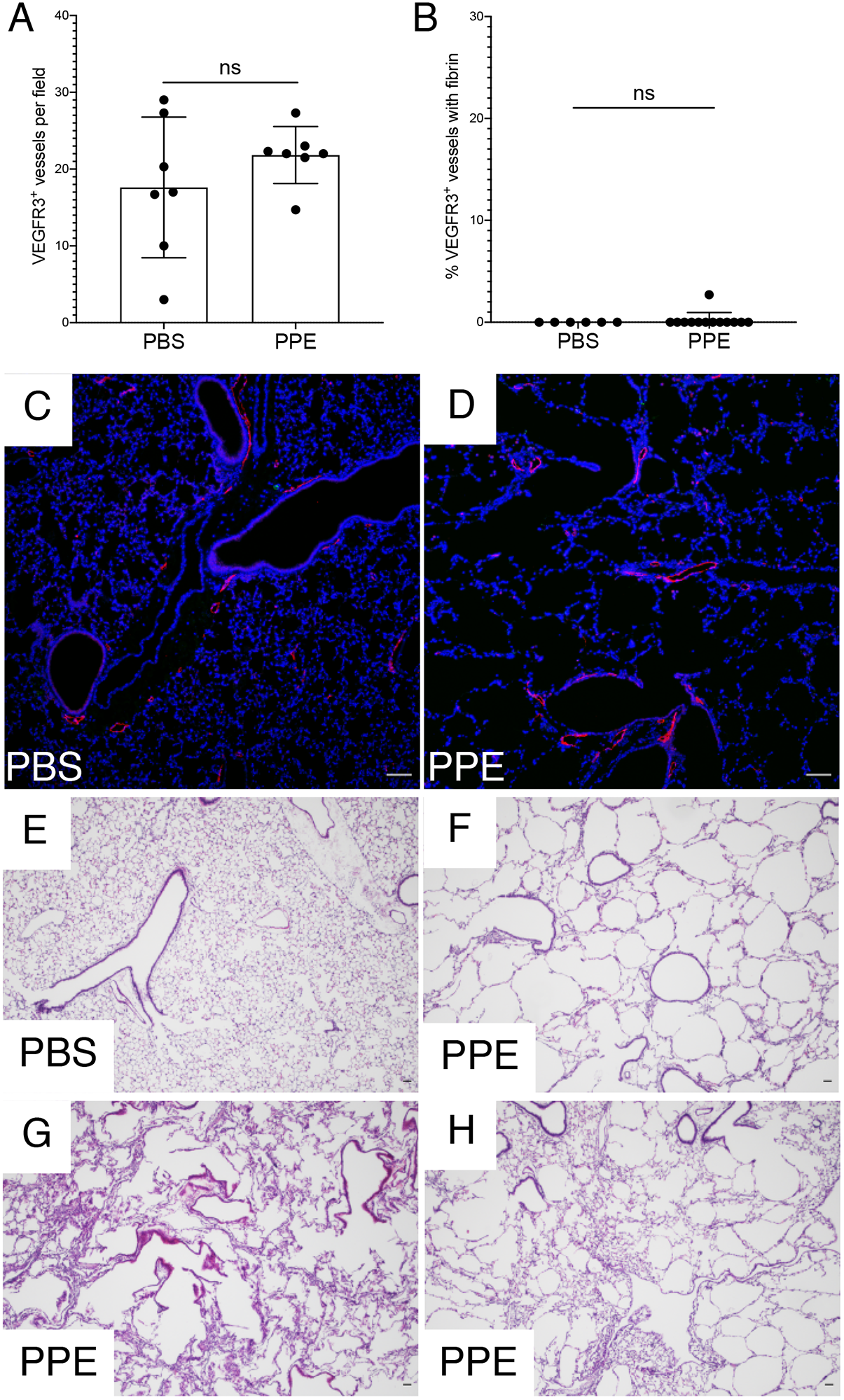
Elastase-induced emphysema does not lead to lung lymphatic thrombosis. (A) Quantification of VEGFR3^+^ lymphatics in lung tissue from mice 21 days after intratracheal porcine pancreatic elastase (PPE) or PBS as a control. (B) Quantification of lymphatic thrombosis, as expressed by percentage of VEGFR3^+^ vessels with luminal fibrin, after PPE or PBS. (C,D) Immunohistochemistry for VEGFR3 (red) and fibrinogen (green) in lung tissue from mice treated with PPE or PBS. (E-H) H&E staining of lung tissue from PPE-treated or PBS-treated mice. At least 5 independent 10x images of lung tissue were used for each mouse tissue sample for quantification. All values expressed as mean ± SEM. *P* value calculated by Student’s *t* test. ns = not significant. Scale bars in E, F = 100µm. All other scale bars = 25µm.

### Cigarette Smoke Extract Causes Increased Lymphatic Endothelial Cell Permeability *in vitro*

The finding of lymphatic thrombosis in human emphysema but not in the elastase model in mice suggested that lymphatic thrombosis may be a result of CS exposure itself, and not the resulting emphysema. To test whether CS has a direct effect on lymphatic integrity, we use used an *in vitro* endothelial cell transport model. Lymphatic endothelial cells (LECs) were seeded on transwell inserts and allowed to reach confluency for 48 hours before being exposed to 1 or 2% (v/v) cigarette smoke extract (CSE) (Figure 3A). After 12 hours, trans-endothelial resistance (TEER) was measured. We found that LEC TEER was decreased with CSE in a dose-dependent manner (Figure 3B). To determine whether the reduction in TEER may be due to changes in LEC cell-cell junctions, we used immunocytochemical staining for the junctional protein VE-Cadherin. We found that LEC cell-cell junctions after CSE appeared jagged and less continuous compared to control LECs (Figure 3C-E). We next tested whether decreased resistance and junctional changes correlated with increased permeability of LECs to bio-inert solutes, specifically 40 kDa and 150 kDa dextran. We found that treatment with CSE led to increased transport of both 150 kDa Dextran and 40 kD dextran across LECs in a time dependent fashion (Figure 3F,G). These studies reveal a direct effect of CSE on LEC permeability.

**Figure 3:**
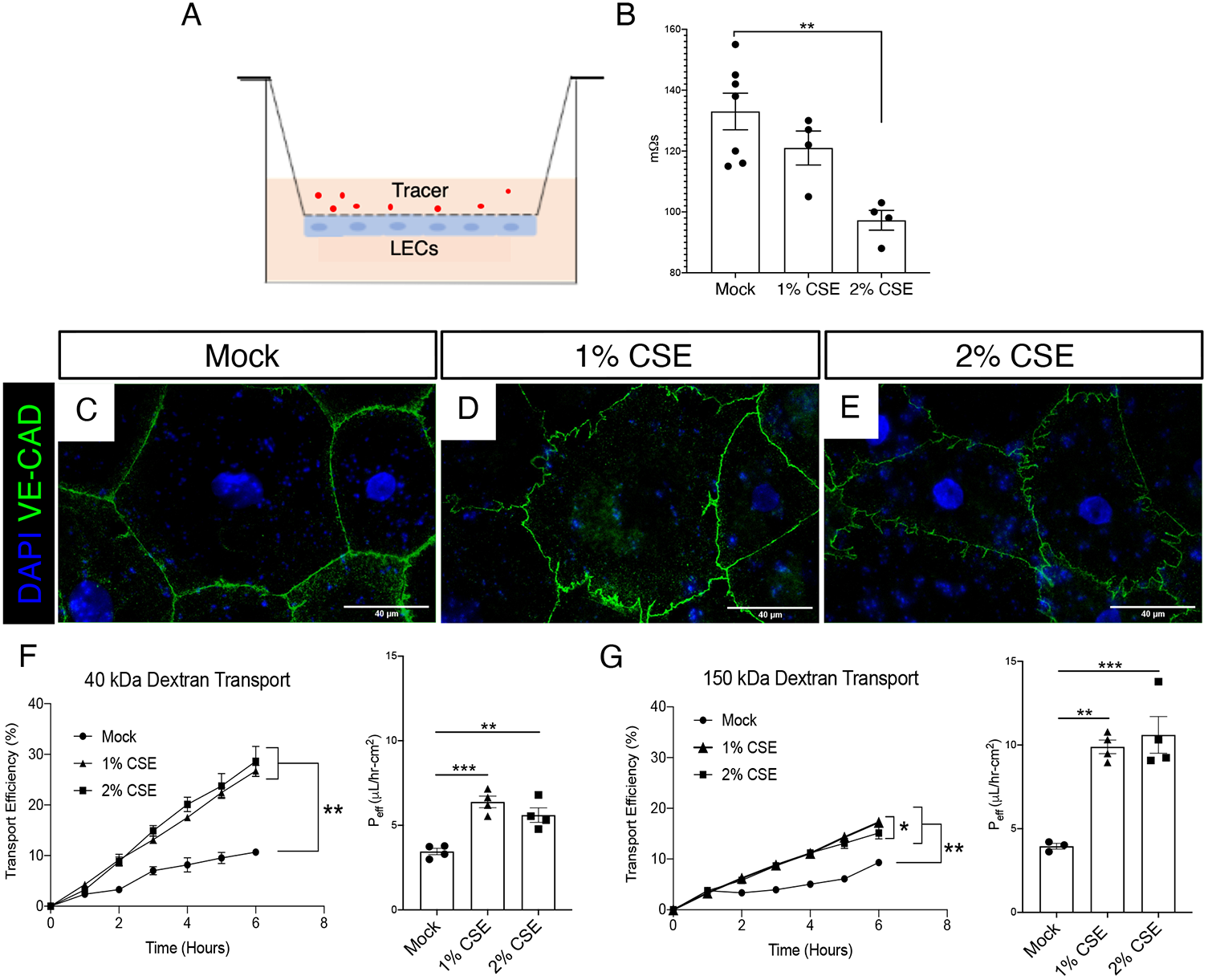
Cigarette smoke extract increases LEC permeability *in vitro*. (A) Schematic of LEC transport model where LECs are seeded on the bottom of a flask and transport of a fluorescent tracer across the monolayer is assessed. (B) Transendothelial electrical resistance (TEER) of a monolayer of LECs after treatment with 1% or 2% CSE for 12 hours. (C-E) Representative fluorescent images of LECs stained for VE-cadherin (VE-CAD, green) after treatment with CSE for 24 hours. (F) Transport efficiency of 40 kDa dextran across LEC monolayer shown over time (left) and as effective permeability (right, P_eff_ µL/h-cm^2^) at 6 hours (n = 3-4). (G) Transport efficiency of 150 kDa dextran across LEC monolayer over time (left) and effective permeability (right, P_eff_ µL/h-cm^2^) at 6 hours (n = 3-4). \**P* < 0.05, \*\**P*< 0.01, \*\*\**P*< 0.001. Scale bars = 40 µm. Microscopy images are representative of at least 4 replicates per group.

### Lung Lymphatic Thrombosis after Cigarette Smoke Exposure in Mice

Given our finding of LEC injury with CSE *in vitro*, we examined the lung lymphatics in mice after CS exposure with a whole body exposure system^15^. Using immunochemistry for the lymphatic marker VEGFR3 which has been shown to be specific for the lung lymphatics in mice^4^, we found no difference in lung lymphatic density after 4 weeks, 8 weeks, or 4 months of smoke exposure compared to identically housed control mice at the same time points (Figure 4A-D). Further analysis revealed fibrin-rich clots in the lymphatics of CS-exposed mice that were not seen in control mice, and in some cases completely obstructed the lumen of the vessel, similar to what we observed in human emphysema (Figure 4E-J). Lymphatic thrombosis was seen after 4 months of CS exposure, but rarely at earlier timepoints (Figure 4J). TLOs are most consistently observed after 6 months or more of CS exposure in mice^16^, and therefore were rarely observed in our samples. However, we identified thrombosed lymphatic vessels that were spatially associated with the sporadic TLOs that were present in the lungs of CS-exposed mice (Figure 4K-M).

**Figure 4:**
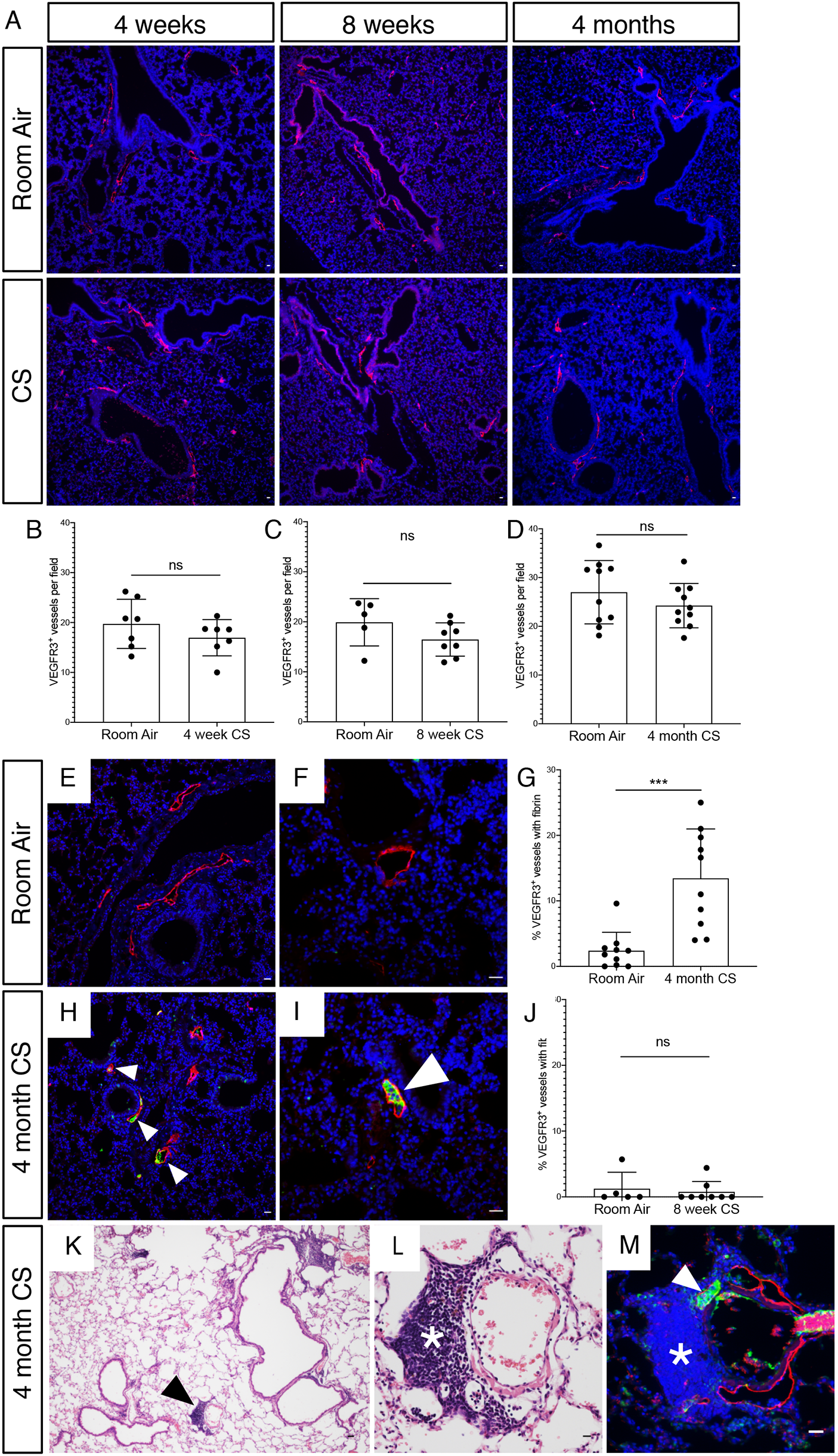
Lymphatic thrombosis after CS exposure in a mouse model of emphysema. Mice were exposed to full body CS and the lungs were harvested for fluorescent immunohistochemical analysis. (A) Representative images of immunohistochemical staining of lung tissue for the mouse lymphatic marker VEGFR3 (red) in mice exposed to CS for 4 weeks, 8 weeks, and 4 months compared to identically housed age matched mice exposed to room air. (B-D) Quantification of lung lymphatic vessel density in CS-exposed and control mice and the indicated time points. (E-J) Analysis of lung lymphatic thrombosis using immunohistochemical staining for VEGFR3^+^ (red) and fibrinogen (green). Thrombosed lymphatics indicated with arrowheads. Quantification of lymphatic thrombosis, as expressed by percentage of VEGFR3^+^ vessels with luminal fibrin after 4 months CS (G) and 8 weeks CS (J). (K,L) H&E of lung tissue after 4 months of smoke exposure, with early TLO indicated with arrowhead (L) and higher magnification of the same area with asterisk in (L). (M) Serial section immunohistochemical staining of TLO seen in (L) for VEGFR3 (red) and fibrinogen (green). Thrombosed lymphatic indicated with arrowhead adjacent to TLO (asterisk). At least 5 independent 10x images of lung tissue were used for quantification. All values expressed as mean ± SEM. *P* value calculated by Student’s *t* test. \*\*\**P* <0.001, ns = not significant. Scale bar in panel K = 100µm. All other scale bars = 25µm.

### Cigarette Smoke Exposure Leads to Decreased Lung Lymphatic Function

We next asked whether there are functional changes in the lung lymphatics in CS-exposed mice that occur prior to emphysema. To assess this, we used a dextran drainage assay, in which fluorescently labeled dextran is delivered to mice intratracheally, and lung lymphatic drainage is quantified by detection of the fluorophore in the mediastinal lymph nodes (mLNs)^4^. Dextran drainage from the lungs to mLNs was significantly decreased in mice after CS exposure, as assessed by both visualization of the mLNs and quantification of mLN fluorescence (Figure 5A-C). Changes in vascular flow are often reflected in endothelial cell morphology, with increased nuclear roundness and changes in nuclear orientation being a hallmark of impaired flow^17^. We used whole mount microscopy in *Prox1-EGFP* lymphatic reporter mice, in which lymphatic endothelial cells express GFP^18^ to further assess the lung lymphatics in CS-exposed mice. We found that lung lymphatic vessels in CS-exposed mice have an altered morphology compared to lung lymphatics in control mice, with rounded lymphatic endothelial cell nuclei that deviated from the axis of flow (Figure 5D,E). These changes were confirmed by quantification of nuclear orientation and length to width ratio (Figure 5F,G).

**Figure 5:**
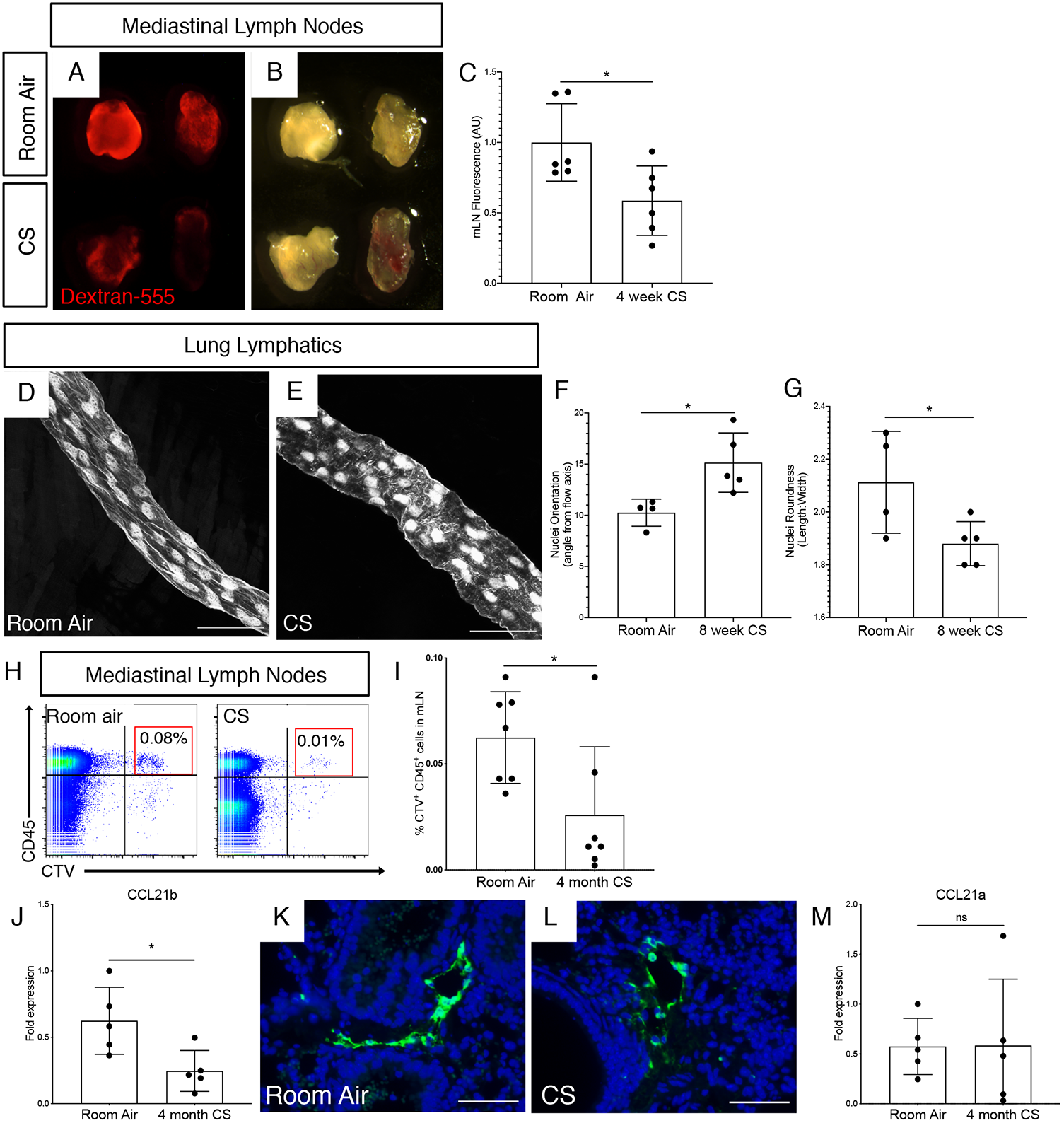
Lung lymphatic dysfunction after CS exposure in mice. Fluorescence (A) and brightfield (B) microscopy of mediastinal lymph nodes (mLN) from mice after 4 weeks CS exposure or room air. mLNs were harvested 60 minutes after intra-tracheal administration of dextran-555 (red). (C) Quantification of dextran-555 uptake to mediastinal lymph nodes by mean fluorescence intensity (AU). (D,E) Whole mount microscopy of lung lymphatics from *Prox1-EGFP* mice exposed to CS (E) or room air (D) for 8 weeks. (F,G) Quantification of nuclear orientation (angle from flow axis) and nuclear roundness (length:width ratio) in lung lymphatic endothelial cells after 8 weeks CS or room air. (H,I) Cell trafficking assay to assess lymphatic leukocyte migration from lungs to draining lymph nodes was performed using intra-tracheal administration of CTV-labeled leukocytes followed by harvest of mLNs for flow cytometry. Identification of CD45^+^ CTV^+^ leukocytes in mLNs of mice exposed to 4 months CS (right) versus room air (left) 48 hours after intra-tracheal administration. (I) Quantification of CTV^+^ leukocytes in mLNs as percent of total cells by flow cytometry. (J) Quantitative PCR for CCL21b using whole lung tissue from mice after 4 months of CS exposure compared to room air. (K,L) Immunohistochemistry for CCL21 (green) using lung tissue from mice after 4 month CS or room air. (M) Quantitative PCR for CCL21a using lung tissue from mice after 4 months of CS exposure compared to room air. Whole mount microscopy images are representative of tissue from at least 4 mice in each group. All values are means ± SEM. *P* value calculated by Student’s *t* test. * *P* <0.05. ns = not significant. Images in A,B shown at 5x magnification. Scale bars in D,E,K,L = 25µm.

A major role of the lung lymphatics is to facilitate migration of immune cells from the lungs to the draining mediastinal lymph nodes^4,19-21^. We tested the effect of CS on lung lymphatic leukocyte trafficking. Leukocytes labelled with cell trace violet (CTV), a fluorescent dye, were administered intratracheally to mice, and their presence in the draining mLNs was detected by flow cytometry^4^. We found significantly decreased lung lymphatic leukocyte trafficking in CS-exposed mice compared to control mice (Figure 5H,I). Furthermore, expression of CCL21b, a cytokine that is expressed by the lung lymphatic capillaries and is critical for leukocyte uptake and migration^22^ was decreased in CS-exposed mice by quantitative PCR as well as immunohistochemical analysis (Figure 5J-L). Interestingly, expression of CCL21a, an isoform in mice that is not specific for the lymphatics and is more broadly expressed in secondary lymphoid tissue^23^ was unchanged in CS-exposed mice by quantitative PCR (Figure 5M). Taken together, these findings indicate decreased lung lymphatic function in CS-exposed mice with thrombosis, decreased drainage, and impaired leukocyte trafficking in these vessels.

### CS Exposure Alters the Composition of Lymph Towards a Prothrombotic and Inflammatory State

We next investigated whether lung lymphatic dysfunction was associated with changes in the composition of lymph in CS-exposed mice. Lymph is a combination of interstitial fluid, products of tissue metabolism, and immune cells, and therefore reflects the physiologic and pathologic signature of the tissue it originated from^1,24,25^. We analyzed the proteomic signature of lymph from CS-exposed mice compared to lymph from control mice exposed to room air. Lymph was harvested from the thoracic duct at a site that is likely to be anatomically enriched for lung drainage (Figure 6A-C). Proteomic analysis revealed a significant number of unique proteins in lymph from CS-exposed mice, as well as changes in the relative abundance of proteins that were expressed in each group (Figure 6D). Pathway analysis of the CS-exposed and control lymph proteome showed upregulation of several proinflammatory pathways, including inflammatory response, cellular damage, cell death and survival, and respiratory disease (Figure 6E). Furthermore, we found upregulation of coagulation pathways in the lymph of CS-exposed mice compared to control, including the extrinsic and intrinsic prothrombin activation pathway (Figure 6F). Upregulation of coagulation pathways included changes in the relative abundance of thrombin, plasmin, Protein C, and Protein S, as well as decreased fibrin and fibrinogen, presumably due to consumption during the formation of clots that we observed in the lymphatics of CS-exposed mice (Figure 7A,B). Interestingly, pathway analysis also showed upregulation of the complement system in the lymph of CS-exposed mice (Figures 6D and 7C). Our findings demonstrate that CS-exposure leads to lung lymphatic dysfunction and changes in the composition of draining lymph that occur prior to the development of emphysema.

**Figure 6:**
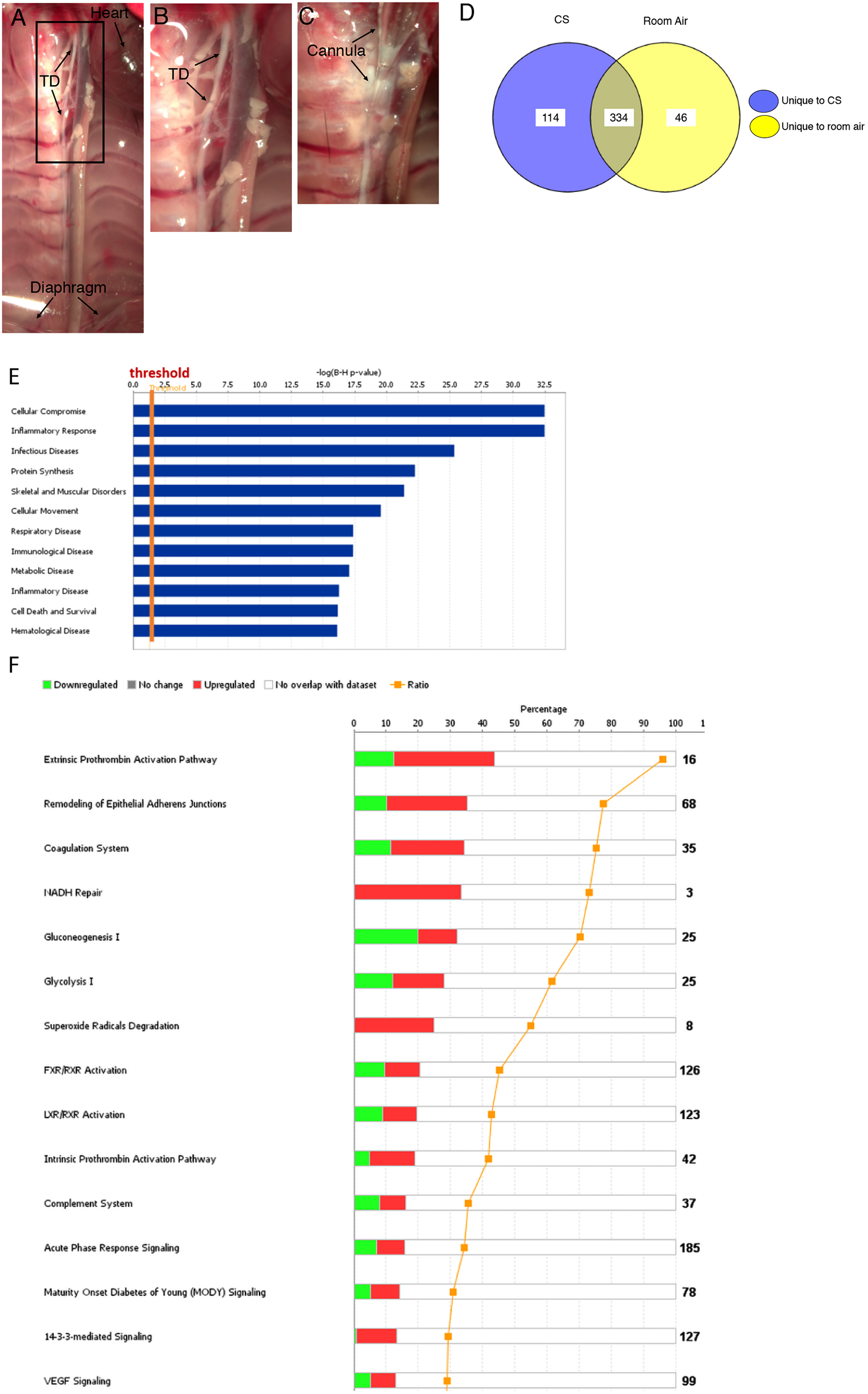
Proteomic analysis of thoracic lymph from cigarette smoke-exposed mice. (A) Brightfield microscopy image demonstrating the thoracic duct (TD) in mice relative to other structures. (B,C) Higher magnification image of region indicated in (A) showing harvest of lymph via cannulation of TD (arrows). (D) Proteomic analysis after 8 weeks of CS identified 114 unique proteins in lymph from CS-exposed mice and 46 unique proteins in lymph from control mice exposed to room air. 334 overlapping proteins were seen in these groups, of which 151 were upregulated and 107 were downregulated in lymph from smoked mice compared to room air. (E,F) IPA of biochemical pathways in the lymph of CS-exposed mice using proteins qualifying for a fold change value of +/- 1.2. Results from 2 independent experiments with a total of n = 9 CS-exposed mice and n = 11 room air control mice. The probability of having a relationship between each IPA indexed biological function and the experimentally determined protein was calculated by right-tailed Fisher’s exact test with the Benjamini-Hochberg Correction. The statistical significance was set to a p-value of <0.05.

**Figure 7:**
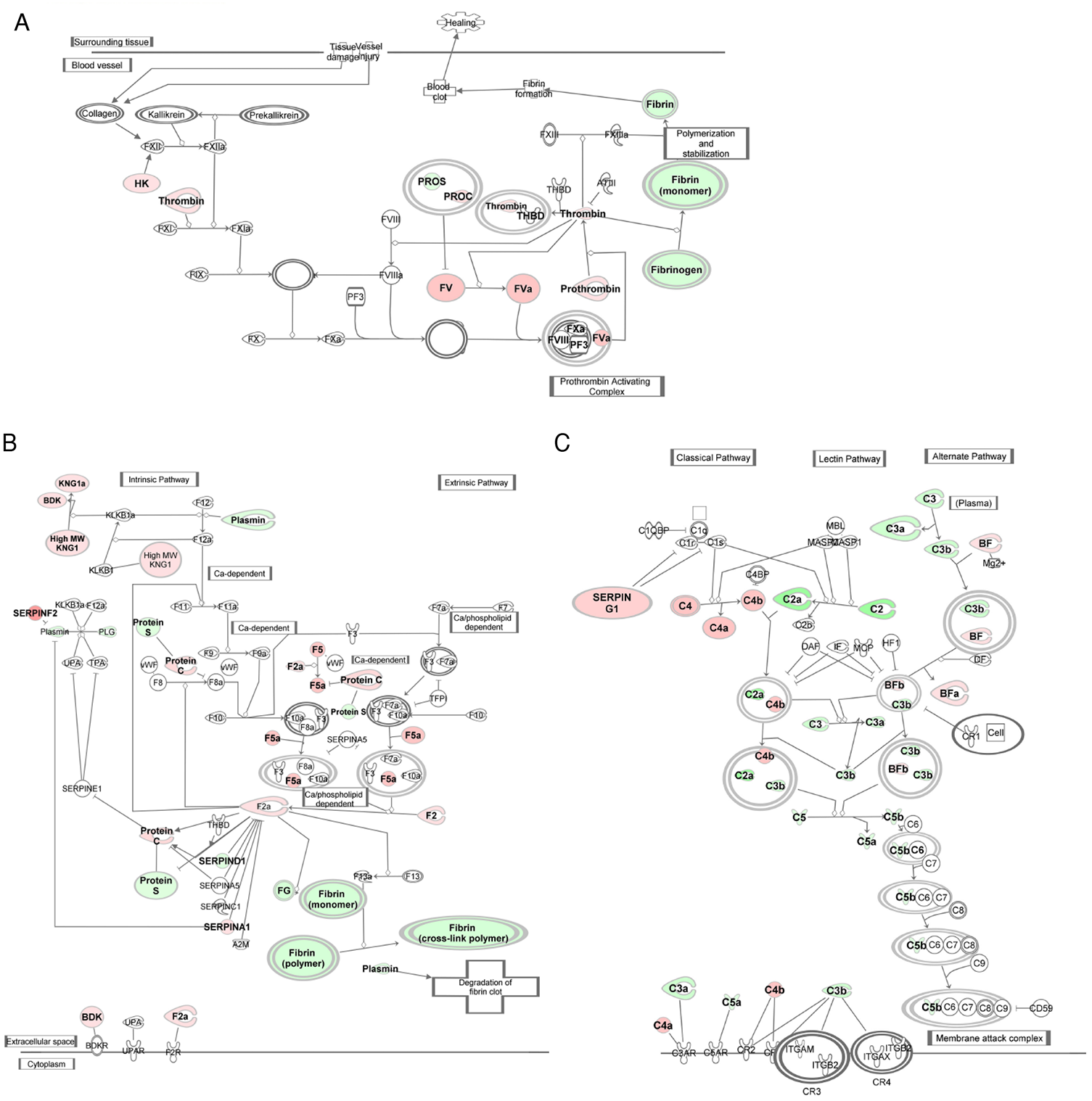
Upregulation of coagulation and inflammation in thoracic lymph from CS-exposed mice. View of IPA-predicted prothrombin activation pathway (A), coagulation system (B), and complement system (C) in lymph from CS-exposed mice compared to room air control mice. Upregulated proteins are shown in red, downregulated proteins are shown in green. Pathway analysis was performed using proteins qualifying for a fold change value of +/- 1.2. Results from 2 independent experiments with a total of n = 9 CS-exposed mice and n = 11 room air control mice.

## Discussion

The lung lymphatics play an important role in lung homeostasis due to their role in fluid drainage and immune cell trafficking. Despite this, lymphatic function in the pathogenesis of chronic lung diseases such as emphysema been less rigorously investigated. Previous studies have documented changes in the density of lung lymphatic vessels in human COPD^12,13^, but whether this reflects changes in lymphatic function is unknown. We have previously shown that lymphatic dysfunction alone results in TLO formation and emphysema in mice. Because TLOs are a prominent feature of CS-induced emphysema^6^, we investigated whether these TLOs are indicative of lymphatic dysfunction and decreased lung lymphatic drainage in this setting. Our studies identified lung lymphatic thrombosis that was significantly increased in patients with emphysema compared to control smokers. Furthermore, lung lymphatic thrombosis was increased in patients with severe emphysema compared to moderate disease. Though not necessarily indicative of impaired lymphatic function, it is probable that lymphatic thrombosis causes impaired lymphatic drainage, as is the case in other settings where lymphatic thrombosis has been observed^26-28^. However, without longitudinal analysis of the human lung tissue, we cannot determine from these data whether lymphatic thrombosis is a predisposing factor associated with severe emphysema, or alternatively, whether severe emphysema itself plays a causal role in subsequent lung lymphatic thrombosis. In addition, though we examined were from patients with a history of smoking, we could not control for active smoking status in these de-identified samples. Due to these limitations, we used mouse models of emphysema to further explore the direct role of CS on lymphatic function.

The most prominent pathologic feature of the lymphatics in CS-exposed mice that we observed in these studies was thrombosis, which was histologically identical to what we observed in human emphysema lung tissue. Functional studies confirmed that lymphatic thrombosis was associated with decreased lung lymphatic drainage and morphologic changes in the lung lymphatic endothelium indicative of impaired lymph flow in mice with CS-exposure. Furthermore, lung lymphatic dysfunction after CS exposure culminated in impaired immune cell trafficking. Lymphatic dysfunction and thrombosis occurred at timepoints that far precede the development of emphysema, which does not occur until 6-8 months of CS exposure in our model^15,29^. The temporal sequence of CS leading to lymphatic dysfunction prior to emphysema was further supported by the finding that severe elastase-induced emphysema alone in the absence of CS exposure was not sufficient to induce lymphatic thrombosis. The absence of lymphatic thrombosis or change in lymphatic density after elastase in mice is striking, given the severe lung injury and emphysema in this model. This suggests a causal role for CS in development lymphatic thrombosis that is independent from emphysema itself. Given the important interplay between the lymphatics and the immune system, CS-induced inflammation may be critical in this process. Taken together, our studies suggest that lymphatic dysfunction and thrombosis are among the initial changes that occur in the lung in response to CS exposure, prior to tissue destruction. This previously unrecognized effect of CS on lung lymphatic function suggests a novel component in the pathogenesis of emphysema.

Lymphatic thrombosis is generally rare and occurs far less frequently than thrombosis in the blood vascular system. This is because despite the presence of fibrinogen and coagulation factors, lymph is generally a hypocoaguable fluid that lacks platelets and has relatively strong fibrinolytic activity^28^. Despite this, lymph does clot in pathologic conditions, and previously reported causes of lymphatic thrombosis include cancer (typically due to external compression and subsequent stasis), infections, heart failure, or chronic edema^26-28,30,31^. To our knowledge, our studies are the first to show lymphatic thrombosis in human emphysema and in response to CS-exposure in a mouse model. CS-induced lymphatic thrombosis may reflect a similar effect of CS on the lymphatic vasculature as in the blood vascular system, where there is well documented endothelial cell injury and coagulation abnormalities^32-34^. Supporting this, we found that cigarette smoke extract causes increased permeability in lymphatic endothelial cells. This increased permeability in lymphatics could promote thrombosis through exposure of tissue factor and activation of the coagulation cascade in a similar manner as is seen in settings of increased blood vascular permeability. Furthermore, our proteomic analysis of lymph from CS-exposed mice showed upregulation of pathways of coagulation and shifts in the relative abundance of proteins that promote clot formation. Thus, the effect of CS on the lymphatic vasculature may involve both direct injury to the lymphatic endothelium as well as changes in the composition of lymph towards a prothrombotic state. In addition, it is conceivable that activated leukocytes that traffic in the lymphatics in the setting of CS exposure may play a role in lymphatic thrombosis and initiate the coagulation cascade. In either scenario, lymphatic permeability, endothelial cell injury, and inflammation in the setting of impaired lymph flow coupled with the prothrombotic effects of CS would fulfill the tenants of ‘Virchow’s triad’ and trigger thrombosis in these vessels.

Here, we have shown that lymphatic thrombosis and dysfunction are early events after CS exposure in mice, and that lung lymphatic thrombosis increases with disease severity in human emphysema. To our knowledge, these studies are the first to demonstrate the functional consequence of CS-exposure on lung lymphatic function. Though not addressed here, our studies raise the possibility that lymphatic dysfunction may either play a role in the pathogenesis in emphysema, or be a marker of disease progression, or both. Given the fundamental role of the lung lymphatics in leukocyte trafficking and regulation of the inflammatory response, a downstream effect of lymphatic dysfunction may be accumulation and activation of immune cells that subsequently cause tissue injury. Importantly, we have previously shown that lymphatic dysfunction alone in mice leads to defects in lung immune cell trafficking, accumulation of these cells, and formation of TLOs^4^. The downstream consequence of lymphatic dysfunction and TLO formation in these mice was in lung injury that resembled emphysema. In light of this, the work presented here raises the question of whether TLOs in human emphysema are also due to lymphatic dysfunction that precedes their formation, and how TLOs formed in this setting contribute to lung injury. The extent to which TLOs due to lymphatic dysfunction play a role in disease progression in emphysema and the mechanism by which this occurs will be the subject of future investigations.

## Materials and Methods

### Human Lung Samples

De-identified human samples were obtained from the NHLBI Lung Tissue Research Consortium (LTRC) biorepository (https://biolincc.nhlbi.nih.gov/studies/ltrc/). These samples were obtained from donor subjects who were planning lung surgery, using tissue that would otherwise be discarded after the lung surgery. Tissue was submitted with a standardized series of questions as well as pulmonary function tests, six-minute walk tests, and chest imaging. We analyzed specimens from control smokers and patients identified as having moderate or very severe COPD, and further restricted our analysis to patients that were identified by the LTRC as having pathologic and radiographic evidence of emphysema.

### Experimental Emphysema

For elastase studies, mice were given 0.25U of porcine pancreatic elastase intratracheally, as previously described^29^. The cigarette smoke exposure studies were performed as previously described^15^. For these studies, *C57Bl/6* or *Prox1-EGFP* reporter mice ^18^ were used. Mice were housed in the Weill Cornell animal facility in 12/12hrs light/dark cycles with ad libitum access to water and food. For all experiments, control and experimental animals were identically housed on the same rack in the animal facility. Both male and female mice were used in both experimental and control groups.

### Functional Assays of Lymphatic drainage and Leukocyte trafficking

Dextran drainage assays and lymphatic leukocyte trafficking using CTV-labelled cells was performed as previously described ^4^. 50ul of 5mg/ml dextran-568 (10,000kD MW, ThermoFisher) was administered to anesthetized, intubated mice via endotracheal catheter. Sixty minutes after administration, the mice were sacrificed for harvest of mediastinal lymph nodes. Lymph nodes were imaged using an Olympus SZX16 dissecting microscope. Quantification of fluorescence intensity was performed using ImageJ.

Cell trafficking experiments were performed as previously described ^4^. Splenocytes were isolated from wild-type mouse spleens and cultured overnight with 100ng/ml LPS and 5ug/ml PHA (Sigma). The cells were then labeled with cell trace violet (CTV, Molecular Probes) according to manufacturer instructions. 1×10^7^ CTV-labeled cells were administered to anesthetized and intubated mice either via endotracheal catheter or intravenously. Lymph nodes or lungs were harvested 48 hours after administration of CTV-labeled cells. Single cell suspensions were stained with the following antibodies: FITC-conjugated anti-CD45 (eBioscience, 11-04551-82), PE-Cy7-conjugated anti-CD11c (BD Bioscience, 561022), and PerCP/Cy5.5-conjugated anti-CD103 (Biolegend, 121415). Flow cytometry was performed using a BD FACSCanto, and analyzed using FlowJo software.

### Whole mount staining

Whole mount staining of lung lymphatics from *Prox1-EGFP* mice was performed according to established protocols ^4^. Tissue from mice carrying the Prox1-EGFP transgene was fixed overnight in 4% PFA at 4°C. For lungs, thick coronal sections were made using a scalpel. Tissue was permeabilized in 0.1% BSA + 0.3% Triton-X in PBS, washed, mounted in Vectashield (Vector) and imaged using a Leica TCS SP8 confocal microscope. Analysis and quantification of nuclear organization and roundness was performed using ImageJ.

### Immunohistochemistry

Fluorescent immunohistochemistry of mouse lung tissue was performed as previously described ^4^. Mice were sacrificed and tissue was perfused with PBS. Lungs were inflated with 4% PFA at constant pressure of 25 cm H_2_O prior to harvest and fixation with 4% PFA overnight. Slides from paraffin-embedded sections were H&E stained or immunostained with antibodies for: VEGFR3 (R&D Systems, AF743) or CCL21 (R&D Systems, AF457), or Fibrinogen (Abcam, ab 227063). Human lung tissue was stained with antibodies for PODOPLANIN (D240, Biolegend, 75782-960) and Fibrinogen (Abcam, ab 227063).

### Quantitative PCR

Quantitative PCR of lung tissue was performed as previously described ^4^. Total RNA from lung tissue was isolated from lung tissue using RNEasy Kit (Qiagen). cDNA was made using Superscript III First-Strand Synthesis System (Invitrogen) following manufacturer instructions. qPCR analysis of gene expression was performed using QuantStudio 6 Real-Time PCR System and SYBR Green PCR Master Mix (Applied Biosystems). Analysis of relative gene expression was carried out using the comparative CT method (ΔCT) using GAPDH as the reference housekeeping gene. Each qPCR reaction was performed in triplicate.

### Lymphatic Permeability Assays

LEC permeability was measured as described, and cigarette smoke extract was generated according to established protocols ^35,36^. LECs were treated with cigarette smoke extract in the culture medium for the indicated amount of time. The underside of 1.0 µM pore size Transwell inserts (Falcon) were coated with 50 µg/mL collagen (Invitrogen). Then human primary LECs were plated at a density of 200,000 cells/cm^2^ on the underside. Models were cultured in EGM-2 (Lonza) at 37°C and 5% CO2 for 48 h to ensure confluence. Transmural fluid flow of 1 µm/s using EGM-2 was introduced for 12 hours to simulate the tissue microenvironment. Then, cigarette smoke extract (CSE) or DMEM (mock) was added to EGM-2 in the top well without added supplemental growth factors. Models were cultured in the new media under flow for two hours before flow was ceased, and 10 µg/mL fluorescently labeled 40 kDa and 150 kDa dextrans (Invitrogen) were added on top. The bottom well was assayed for fluorescence every hour for up to 12 h. Fluorescence intensity was measured using a plate reader (Tecan) and the amount of tracer transported was calculated using a standard curve. Effective permeability was estimated using the equation:

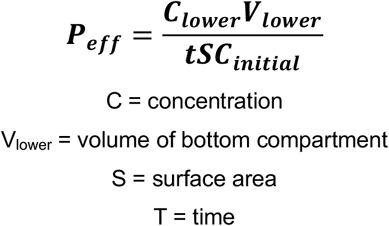

LEC monolayer was confirmed using immunofluorescence staining and trans endothelial and epithelial resistance (TEER, Millipore Sigma) was measured 12 hours after CSE treatments.

### Immunofluorescence staining of LECs used for Lymphatic Transport Model

Cells were fixed in 2% PFA for 15 minutes at room temperature and incubated with mouse anti-human VE-Cadherin (BD Sciences) at 4°C overnight as previously described ^36^. Secondary antibodies labeled with Alexa Fluor® 488 were used for detection. Slides were mounted using DAPI-containing Vectashield (Vector Laboratories) and imaged using a Zeiss Axio Observer. Image processing was performed using FIJI.

### Lymph Harvest and Proteomic Analysis

Thoracic lymph was harvested from mice and samples were processed for proteomic analysis as previously described ^37^ and detailed in the online methods supplement.

### Statistics

Data are expressed as the mean ± SEM, and the numbers of samples per condition are indicated in the figure legends. Statistical significance was determined by unpaired, 2-tailed Student’s t-test or ANOVA using GraphPad Prism software. P values of less than 0.05 were considered statistically significant. Quantification of lymphatics in lung tissue was performed using at least 5 randomly captured 10x images of VEGFR3 staining per mouse. Lymphatic thrombosis was quantified as the number of VEGFR3^+^ lymphatic vessels with fibrin present in the lumen, as a percentage of total VEGFR3^+^ vessels in the image. For human samples, number of lymphatics was calculated as the number of podoplanin^+^ vessels per 10x image. At least 3 randomly captured 10x images were used per sample. Lymphatic thrombosis was calculated as podoplanin^+^ lymphatic vessels with fibrin in the lumen, expressed as a percentage of total podoplanin^+^ vessels. TLOs were defined as discrete lymphocyte-dense accumulations on H&E-stained sections. Number of TLOs was quantified using 10x images representing the entirety of the lung tissue section for each patient sample.

### Study approval

All animal experiments were approved by the IACUC of Weill Cornell Medicine and performed in accordance with relevant guidelines and regulations. Reporting in the manuscript follows the recommendations in the ARRIVE guidelines.

## Acknowledgements

The authors wish to the thank the Microscopy and Image Analysis Core Facility at Weill Cornell Medicine, the Histology Core Facility at Weill Cornell Medicine, and the Proteomics and Macromolecular Crystallography Shared Resource at the Herbert Irving Comprehensive Cancer Center of Columbia University Medical Center.

## Author Contributions

Conception and design of work: H.O.R., K.M., L.M., L.S., A.M.K.C, M.L.K. Acquisition, analysis, and interpretation of data: B.D.S., K.K., S.T., A.C.R., S.Z., S.Q., C.C.C., Z.K., J.Y., K.M., J.M., J.D’A, H.O.R. Manuscript preparation and editing: B.D.S. and H.O.R. Final manuscript review: All authors.

Supported by the NHLBI K01HL145365 (to H.O.R), the Robert Wood Johnson Foundation (to H.O.R.), NIH AI146180 and NIH AI137198 (to L.S.), NIGMS MIRA 1R35GM142835-01 (to K.M.), and AHA Predoctoral Fellowship 834524 (to J.M.)

The author(s) declare no competing interests.

## Supplementary Methods

### Lymph Harvest and Proteomic Analysis

#### Chemicals and other reagents

Acetic acid (ULC/MS grade), acetonitrile, formic acid, methanol, trifluoroacetic acid (TFA), and ULC/MS-grade water (for nano-LC analysis, 99% purity) were purchased from Fisher Scientific. TCEP-HCl, iodoacetamide, ammonium bicarbonate, glycine, urea, thiourea, KCl, KH_2_PO_4_, H_3_PO_4_, and Na_2_CO_3_ were of the highest grade available from Sigma-Aldrich Millipore. Porcine trypsin (20 mg, specific activity >5,000 units/mg sequencing grade modified), Lys-C (sequencing grade, 10 mg) and Glu-C, sequencing grade (10 mg) were purchased from Promega (Madison, WI). All solutions were prepared using MilliQ water purified by an Elix 3 UV Water Purification System (Millipore, Billerica, USA) and filtered through 0.2 mm pore membrane sterile filter units (Steritop−, Millipore). All methods were performed in accordance with the relevant guidelines and regulations. Total protein quantitation was performed using the Micro BCA− Protein Assay Kit, (cat # 23235 from Thermofisher Scientific). Amicon Ultra-0.5 ml centrifugal filters (Ultracel-10K, cat#UFC501024) were purchased from Millipore-Sigma.

#### Extraction of endogenous peptides (peptidome) from lymph sample

Equal amounts of 50-100 ug of total protein from the lymph collected from mice exposed CS-exposed mice (n = 9) and room air controls (n = 11) were equilibrated in 0.4 ml of sterile PBS buffer supplemented with a cocktail of protease inhibitors (Roche). Peptides were extracted using 0.2% TFA. Samples were then filtrated through a 10,000-Da cutoff filter device (Amicon) at 10°C for 30 minutes, desalted using pepClean C-18 spin columns (Pierce), and eluted with 70% acetonitrile containing 0.1% TFA for further nanoLC/MS/MS analysis.

#### Processing of lymph samples for proteomics analysis

50-100 mg from each lymph were reduced with 25 mM TCEP-HCl (Thermo Scientific) in 50 mM ammonium bicarbonate (ABC) buffer, containing 8 M urea at pH 8.5 for 45 minutes at RT followed by alkylation with 100 mM iodoacetamide for 50 minutes in the dark at RT. The protein solutions were transferred on microcon-10kDa centrifugal filter units with Ultracel-10 membrane (catalog# MRCPRT010) from Millipore Sigma and washed with 50 mM ammonium bicarbonate buffer five times at 9000xg in a microcentrifuge, for 10 minutes each step. The reduced and alkylated samples were resuspended in 100 ml of 50 mM ABC buffer, pH 8.9 (urea <2M) and digestion was carried out at 37°C overnight (12 hours) using a combination of three enzymes: trypsin/LysC at 20:1 protein: enzyme ratio and GluC at 10:1 protein: enzyme ratio. The digestion was quenched with 0.5% acetonitrile and 1.5% formic acid. Processed peptides were then extracted through a 10-kDa MWCO (molecular weight cut-off) using 10kDa centrifugal filter units by spinning at 10,000xg for 15 minutes in a 20:1 microcentrifuge. The final peptide mixture, extracted from all enzymatic digestions, was desalted on C18 Prep clean columns (EMD Millipore) and reconstituted in 25 µl 5% acetonitrile containing 0.1% (v/v) trifluoroacetic acid for further nanoLC/MS/MS analysis.

Equal aliquots (ug) of endogenous peptides (MW<10 kDa) and/or tryptic peptides were analyzed in replicates (n=22 biological replicates for each sample set) by nano-LC-MS/MS using a combination of data dependent and independent analyses (DDA and DIA, respectively). We used a Thermo Scientific− Orbitrap Fusion− Tribrid− mass spectrometer and applied a protocol developed and published previously (Clement CC et al., Immunity 2021).

Briefly, desalted peptides were injected onto an EASY-Spray PepMap RSLC C18 50 cm x 75 μm column (Thermo Scientific), which was coupled to the mass spectrometer. Peptides were eluted with a non-linear 180 min gradient of 5-30% buffer B (0.1% (v/v) formic acid, 100% acetonitrile) at a flow rate of 250 nL/min. The column temperature was maintained at a constant 50 °C during all experiments. For DIA analysis, survey scans of peptide precursors were performed from 350-1200 *m/z* at 120K FWHM resolution (at 200 *m/z*) with a 1 × 10^6^ ion count target and a maximum injection time of 60 ms. The instrument was set to run in top speed mode with 3s cycles for the survey and the MS/MS scans. After a survey scan, 26 m/z DIA segments were acquired from 200-2000 *m/z* at 60K FWHM resolution (at 200 *m/z*) with a 1 × 10^6^ ion count target and a maximum injection time of 118 ms. HCD fragmentation was applied with 27% collision energy and resulting fragments were detected using the rapid scan rate in the Orbitrap. The spectra were recorded in profile mode.

#### DDA nano-LC/MS/MS for generation of spectral libraries

The sample specific spectral library (SSL) was generated by pooling 1/10 aliquots from each biological sample and DDA method for peptide MS/MS analysis. Survey scans of peptide precursors were performed from 400 −1500 *m/z* at 120K FWHM resolution (at 200 *m/z*) with a 4 × 10^5^ ion count target and a maximum injection time of 50 ms. The instrument was set to run in top speed mode with 3 s cycles for the survey and the MS/MS scans. After a survey scan, tandem MS was performed on the most abundant precursors exhibiting a charge state from 2 to 6 of greater than 5 × 10^3^ intensity by isolating them in the quadrupole at 1.6 Th. CID fragmentation was applied with 35% collision energy and resulting fragments were detected using the rapid scan rate in the ion trap. The AGC target for MS/MS was set to 1 × 10^4^ and the maximum injection time limited to 35 ms. The dynamic exclusion was set to 60 s with a 10-ppm mass tolerance around the precursor and its isotopes. Monoisotopic precursor selection was enabled. The remaining half of each sample was run using DIA method as described above.

#### Generation of Spectral Libraries

To generate the spectral libraries, the acquired DDA raw files corresponding to the pooled samples from the biological replicates (aliquots of 1/10 from each CS-exposed and room air control sample) were searched with PEAKS X+ and then filtered with Scaffold software (version 4.6.2). The enzyme restriction was set up as “*no enzyme*” option in PEAKS X+ to fit the endogenously processed peptides. Then, the spectral library was exported as a *“*.*blib”* file using the built-in available option from the Scaffold software.

An independent analysis of the MS/MS DDA raw files was performed using the MSFragger (The Nesvizhskii Lab, 1301 Catherine, 4237 Medical Science I, Ann Arbor, MI 48109: version 3.2). MSFragger was set up to search a reverse concatenated uniprot-filtered-organism Mus musculus (mouse) database (April 2021, 36,902 entries) assuming the digestion enzyme trypsin for proteomics samples. MSFragger was searched with a fragment ion mass tolerance of 20 PPM and a parent ion tolerance of 20 PPM. Deamidated of asparagine and glutamine, oxidation of histidine, methionine, and tryptophan and carbamidomethyl of cysteine were specified in MSFragger as fixed modifications. Scaffold (version Scaffold_5.0.1, Proteome Software Inc., Portland, OR) was used to validate MS/MS based peptide and protein identifications. Peptide identifications were accepted if they could be established at greater than 95.0% probability by the Scaffold Local FDR algorithm. Protein identifications were accepted if they could be established at greater than 95.0% probability and contained at least 1 identified peptide. Protein probabilities were assigned by the Protein Prophet algorithm (Nesvizhskii, Al et al Anal. Chem. 2003;75(17):4646-58). Proteins that contained similar peptides and could not be differentiated based on MS/MS analysis alone were grouped to satisfy the principles of parsimony. Then, the spectral library was exported as a “.blib” file using the built-in available option from the Scaffold software.

#### DIA Analysis of peptidomes and proteomics data

DIA data were analyzed using Scaffold DIA (1.2.1) (Proteome Software Inc., Portland) which had the raw data files converted to mzML format using ProteoWizard (3.0.11748). The analytic samples were aligned based on retention times and individually searched against “DDA.blib spectral library” with a peptide mass tolerance of 10 to 15 ppm and a fragment mass tolerance of 15 to 50.0 ppm. Variable modifications were imported from the DDA based spectral library as follow: methionine, lysine, proline, arginine, cysteine, and asparagine oxidations (+15.99 on CKMNPR), deamidation of asparagine and glutamine (NQ-0.98) and pyro-Glu from glutamine (Q-18.01 N-term). The *“no enzyme*” option was used in Scaffold DIA with variable allowed 8-12 missed cleavage site(s) for peptidomics analysis. Only peptides with charges in the range [2-8] and length in the range [5-25] were exported for further quantitation. For the proteomics analysis, “trypsin” restriction was used for the enzyme digestion and one “allowed missed cleavage” Peptides identified in each sample were filtered by Percolator (3.01. nightly-13-655e4c7-dirty) to achieve a maximum FDR between 0.01-0.05. Individual search results were combined, and peptide identifications were assigned posterior error probabilities and re-filtered to FDR thresholds of 0.01-0.05 by Percolator (3.01. nightly-13-655e4c7-dirty). Peptide quantification was performed by Encyclopedia (0.7.2). For each peptide, the 5 highest quality fragment ions were selected for quantitation. The intensities for the proteins were calculated and normalized by summation of the peptide intensities using the Scaffold DIA’s built-in normalization algorithm.

#### Independent analysis of DIA data files from peptidomes and proteomics with PEAKS X+/pro

Raw files from each biological replicate were filtered using the DIA built-in option in PEAKS X+ /Pro(Bioinformatics Solutions, Waterloo, Canada), de novo sequenced, and assigned with protein ID using by searching against the mouse Swiss-Prot database (April 2021; 36,902 entries) and the following search parameters trypsin/LysC/Glu-C, as restriction enzymes in the case of proteomics data set and “*no enzyme*” for searching the peptidomics dataset; two allowed missed cleaves at one peptide end was applied for tryptic peptides. The parent mass tolerance was set to 13 ppm using monoisotopic mass, and fragment ion mass tolerance was set to 0.03 Da. Carbamidomethyl cysteine (+57.0215 on C) was specified as a fixed modification. Methionine, lysine, proline, arginine, cysteine, and asparagine oxidations (+15.99 on CKMNPR), deamidation of asparagine and glutamine (NQ-0.98), and pyro-Glu from glutamine (Q-18.01 N-term) were set as variable modifications. We also performed an additional analysis of peptidomes and proteomics DIA files using the “*Spectral Libraries*” search option enabled by the PEAKS X/Pro, using the spectral libraries generated from DDA raw files as described above.

Data were validated using the false discovery rate (FDR) method built in PEAKS X+, and protein identifications were accepted with a confidence score (−10lgP) >15 for peptides and (−10lgP) >15 for proteins; a minimum of one peptide per protein was allowed after data were filtered for <5% FDR for endogenous peptides and FDR<1% for tryptic peptides; and <3% FDR for proteins identifications (P < 0.05).

